# High cRel-expressing germinal center B cells favor precursor plasma cell generation

**DOI:** 10.64898/2026.04.26.720943

**Authors:** Sukanya Roy, Tracy Tabib, Bipasha Mazumder, Ashok Kumar, Easton Van De Walle, Saurabh Kumar, Jishnu Das, Koushik Roy

**Affiliations:** Division of Microbiology and Immunology, Department of Pathology, University of Utah, Salt Lake City, UT 84112; Center for Systems Immunology, Department of Immunology, University of Pittsburgh, Pittsburgh, PA 15260; Huntsman Cancer Institute, University of Utah, Salt Lake City, UT 84112

**Keywords:** Germinal Center B Cell, GC B cell selection, Precursor Plasma Cell, NFκB, cRel, transcription regulation

## Abstract

Germinal centers (GCs) are the site of antibody affinity maturation by the process of somatic hypermutation (SHM) and produce long-lived plasma cells (PCs). GC B cells circulate between two distinct zones: the light zone (LZ) and the dark zone (DZ). It has been demonstrated that the transcription factor NFκB cRel is required for B cell proliferation and GC B cell maintenance. cRel-deleted GC B cells show reduced SHM and affinity maturation. In contrast, transgenic overexpression of cRel does not affect SHM and results in little or no increase in affinity maturation. Therefore, the function of cRel in regulating SHM and GC B cell-derived PC generation remains unknown. To understand the function of cRel in GC B cell selection, we have used cRel fluorescence reporter mice, which provide insight into cRel function in GC B cells at the level of natural expression variation. We found that about 6-9% of LZ cells increased cRel expression compared to DZ cells, and high cRel-expressing LZ cells showed increased SHM and increased expression of IRF4 and cyclin D2, though cMyc expression remained similar. Combining single-cell RNA-Seq and flow cytometry, we revealed that high cRel-expressing LZ cells are enriched for precursor PCs (pre-PCs), not precursor memory B cells (pre-MBCs). Our findings provide insight into the physiologic function of cRel in regulating GC B cell output.

## Introduction

GCs are transient structures formed in response to antigen encounter by immunization, infection and autoimmunity. GCs are the site of B cell proliferation coupled with SHM of the B cell receptors (BCRs) to produce high-affinity antibodies (1-4). GC B cells circulate between two distinct zones: the LZ and the DZ. SHM occurs in the DZ, and SHM is a random process that could produce BCR with alter affinity or produce damage and nonfunctional BCRs. B cells with damage and nonfunctional BCRs die in the DZ, and the remaining live DZ cells cycle to the LZ for positive selection. LZ cells are selected based on their BCR affinity to differentiate into PCs and MBCs in LZ. Following selection in the LZ, GC B cells can also reenter the DZ for additional rounds of division and SHM (1-4). LZ cells that differentiate into PCs and MBCs are rare, yet well-designed recent studies have identified precursor cells for both, as well as transcription factors and cell-cycle activity controlling GC-derived MBCs and PCs (5-10). *Hhex* and *Tle3* are key transcriptional regulators of precursor MBCs (pre-MBCs) differentiation, and pre-MBCs show enhanced expression of cell-survival regulator Bcl2 (2, 7-9). IRF4 and Bcl6 are key regulators of precursor PCs (pre-PCs) differentiation, and pre-PCs show enhanced expression of the cell-cycle regulator Cyclin D2 (6). Higher expression of the cell-cycle regulator cMyc characterizes positively selected GC B cells (11-13). Recent studies show that high cMyc LZ cells differentiate into PCs and MBCs and reenter DZ (14, 15). cMyc expression in B cells is controlled by cRel (16, 17).

The transcription factor NFκB cRel is associated with many human autoimmune diseases, and GC B cells play a central role in the development of autoimmunity (18-20). Gain of function of the *Rel* gene, which encodes cRel protein, is associated with human B cell lymphomas, and many B cell lymphoma originate from GC B cells (21, 22). cRel in GC B cells is activated by CD40 signaling alone and in combination with BCR signaling (13). It has been demonstrated that cRel is required for GC B cell maintenance, and cRel deletion in GC B cells reduces SHM and affinity maturation (23). Transgenic overexpression of cRel in GC B cells enhances GC B cell formation, though surprisingly it does not affect SHM and, if any, reduces affinity maturation (19). Hence, the function of cRel in SHM and positive selection remains unknown. Studies on transcription factors, SOX9 and GFI1B, have shown differential functions under physiologic expression and genetic perturbation (24, 25).To elucidate the function of cRel heterogeneity in GC B cell selection at physiologic expression levels, we have utilized recently generated cRel fluorescence reporter mice (mTFP1-cRel). We found that cRel expression is higher in GC B cells compared to MBCs and other B cell types. Further analysis showed that approximately 6-9% of LZ cells had higher cRel expression than DZ cells, and the highest cRel-expressing LZ cells have increased SHM and higher expression of IRF4 and Cyclin D2, while cMyc expression remains similar between high- and low-cRel LZ cells. Single-cell RNA-seq shows that pre-PCs exhibit increased cMyc expression, as previously reported (14, 15). Combining single-cell RNASeq analysis and flow cytometry, we show that cRel high LZ cells are enriched for pre-PCs rather than for pre-MBCs. Overall, our study suggests that at the physiologic cRel expression level, high cRel-expressing LZ cells enhance SHM and are enriched for pre-PCs.

## Results and discussion

### cRel expression is higher in GC B cells than in MBCs

To determine the function of physiologic cRel expression in the GC B cells, we utilized cRel fluorescence fusion reporter mice. In this mouse strain, cRel is expressed as a knock-in with monomeric teal fluorescent protein 1 (mTFP1) and expressed as a fusion protein with cRel as described previously and defined here as mTFP1-cRel (cRel^mTFP1^) (17). To test the effect of mTFP1 fusion to cRel on GC B cell formation, we immunized C57BL/6J (WT) and cRel^mTFP1^ mice and measured GC B cell formation. We found that WT and cRel^mTFP1^ mice generated similar proportions of GC B cells (WT and cRel^mTFP1^ mean GC B cells proportion: 10.2% and 10.6%, respectively) (Fig. 1A-1C and Fig. S1A). Further, we found that the number of GC B cells in WT and cRel^mTFP1^ remained similar (Fig. 1D). We measured cRel expression in GC B cells and other B cell types (defined here as non-GC B cells) and found that cRel expression was higher in GC B cells than in non-GC B cells in cRel^mTFP1^ mice, with 26.7% mean increase in cRel expression in GC B cells (Fig. 1E-1F). To confirm increased cRel expression in GC B cells compared to non-GC B cells, we measured intracellular cRel expression with a cRel monoclonal antibody in WT mice. We found that cRel expression was higher in GC B cells than in non-GC B cells, with 62% mean increase in cRel expression in GC B cells (Fig. 1G-1H). The slight differences in mean cRel expression changes between GC and non-GC B cells may be attributed to differences in flow cytometry methods used for fixed and unfixed cells. The observation of increased cRel in GC B cells compared to non-GC B cells is consistent with prior findings (19, 26, 27).

**Figure 1:**
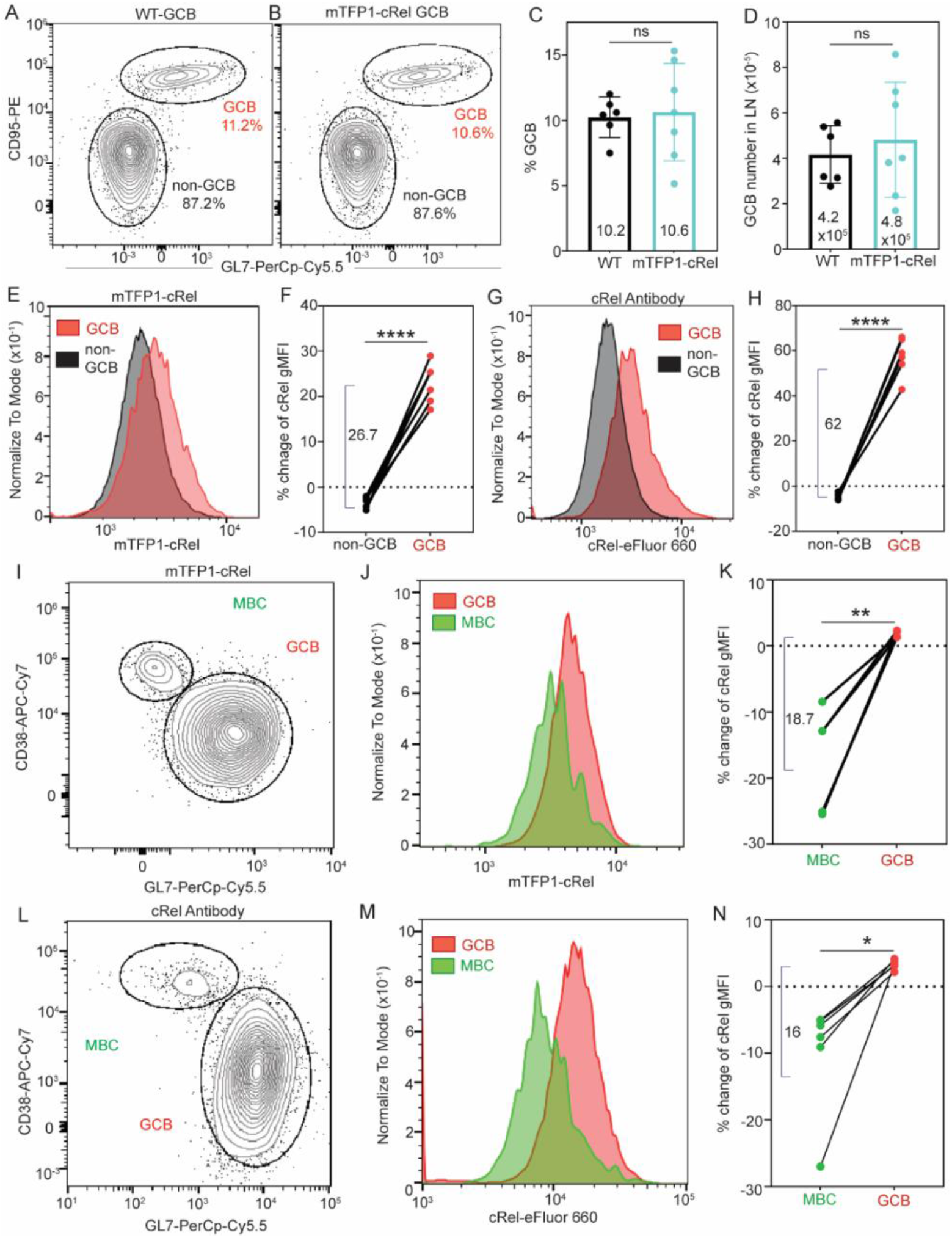
Increased cRel expression in GC B cells compared to MBCs. WT and cRel^mTFP1^ mice were immunized, and the generation of GC B cells (B220+CD95+GL7+) and non-GC B cells (B220+CD95-GL7-) was measured. **(A)** Flow cytometry plot of GC B cells and non-GC B cells proportion in WT mice. **(B)** Flow cytometry plot of GC B cells and non-GC B cells proportion in cRel^mTFP1^ mice. **(C)** Proportion of GC B cells within the total B cell population in WT and cRel^mTFP1^ mice. The mean proportion of GC B cells is indicated within the bar (WT mean 10.2 and cRel^mTFP1^ mean 10.6). **(D)** The total number of GC B cells per lymph node (LN) in WT and cRel^mTFP1^ mice. The total number of GC B cells is indicated within the bar (WT mean 4.2×10^5^ and cRel^mTFP1^ mean 4.8×10^5^). **(E)** Histogram of cRel expression in GC B cells and non-GC B cells in cRel^mTFP1^ mice. **(F)** Proportion changes of cRel gMFI (geometric mean fluorescence intensity) in GC B cells and non-GC B cells from total B cells in cRel^mTFP1^ mice. gMFI represents cRel expression. 26.7 is the mean % increase in cRel expression in GC B cells compared to non-GC B cells. The proportion change of cRel expression is calculated as follows: 100x(X-Y)/Y. X represents the gMFI of GC B cells or non-GC B cells, and Y represents the gMFI of total B cells, which includes both GC B cells and non-GC B cells. **(G)** Histogram of cRel expression in GC B cells and non-GC B cells in WT mice stained with cRel antibody. **(H)** Proportion changes of cRel gMFI in GC B cells and non-GC B cells from total B cells in WT mice stained with cRel antibody. 62 is the mean % increase in cRel expression in GC B cells compared to non-GC B cells. **(I)** Flow cytometry plot of GC B cells (B220+CD19+CD3e-NK1.1-IgD^low^CD38-GL7+) and MBCs (B220+CD19+CD3e-NK1.1-IgD^low^CD38+GL7-) in cRel^mTFP1^ mice. **(J)** Histogram of cRel expression in GC B cells and MBCs in cRel^mTFP1^ mice. **(K)** Proportion changes of cRel gMFI in GC B cells and MBCs from IgD^low^ B cells in cRel^mTFP1^ mice. gMFI represents cRel expression. 18.7 is the mean % increase in cRel expression in GC B cells compared to MBCs. The proportion change of cRel expression is calculated as follows: 100x(X-Y)/Y. X represents the gMFI of GC B cells or MBCs, and Y represents the gMFI of IgD^low^ B cells, which includes both GC B cells and MBCs. **(L)** Flow cytometry plot of GC B cells and MBCs in WT mice stained with cRel antibody. **(M)** Histogram of cRel expression in GC B cells and MBCs in WT mice. **(N)** Proportion changes of cRel gMFI in GC B cells and MBCs from IgD^low^ B cells in WT mice. gMFI represents cRel expression. 16 is the mean % increase in cRel expression in GC B cells compared to MBCs. One dot represents one mouse. For **(C)** and **(D)**, an unpaired Student’s t-test was performed. For **(F), (H), (K)**, and **(N)**, a paired Student’s t-test was performed. ns: not significant, *p < 0.05, **p < 0.01, ***p < 0.001, and ****p < 0.0001.

We distinguished MBCs from non-GC B cells and investigated whether cRel expression in MBCs was similar to or distinct from that in GC B cells. We found that cRel expression in MBCs is lower than that of GC B cells in cRel^mTFP1^ mice, with 18.7% mean decrease in cRel expression in MBCs (Fig. 1I-1K and Fig. S1B). To confirm reduced cRel expression in MBCs compared to GC B cells, we measured intracellular cRel expression with the cRel monoclonal antibody in WT mice. We found that cRel expression is reduced in MBCs compared to GC B cells in WT, with 16% mean decrease in cRel expression in MBCs (Fig. 1L-1N), thus confirming reduced cRel expression with both fluorescence reporter and intracellular staining. In conclusion, we show that GC B cells express higher levels of cRel than MBCs and other B cell types.

### High cRel-expressing LZ GC B cells show increased SHM

Previous studies have shown that cMyc expression is higher in a small fraction of LZ cells than in DZ cells, and high cMyc-expressing LZ cells are positively selected and preferentially enter the DZ for additional SHM (4, 11, 12). It has been shown that cRel-deleted B cells fail to induce cMyc expression upon mitogenic stimulation (16). Recently, we have demonstrated that cMyc expression correlates with cRel activation in mature B cells (17). Therefore, we hypothesized that cRel expression is higher in LZ cells than in DZ cells, and high cRel expression is associated with Myc expression in GC B cells. To test this hypothesis, we measured cRel expression in LZ and DZ cells. We found that cRel expression was higher in LZ cells from cRel^mTFP1^ immunized mice, with 29% mean increase in cRel expression in LZ cells relative to DZ cells (Fig. 2A-2C). We quantified the proportion of LZ cells that increase in cRel expression and found that ∼8.4% LZ cells express higher cRel than DZ cells (Fig. 2D). To confirm increased cRel expression in LZ cells compared to DZ cells, we measured intracellular cRel expression with cRel monoclonal antibody in WT mice. We found that cRel expression was higher in LZ cells than in DZ cells in WT, with a 34% mean increase in cRel expression in LZ cells relative to DZ cells (Fig. 2E-2G). We quantified the proportion of LZ cells that increase in cRel expression and found that ∼5.2% LZ cells expressed higher cRel than DZ cells (Fig. 2H). Hence, confirming increased cRel expression using both a fluorescence reporter and intracellular staining. The observation of increased cRel in GC B cells is consistent with a recent finding that cRel expression is higher in LZ cells than in DZ cells (19).

**Figure 2:**
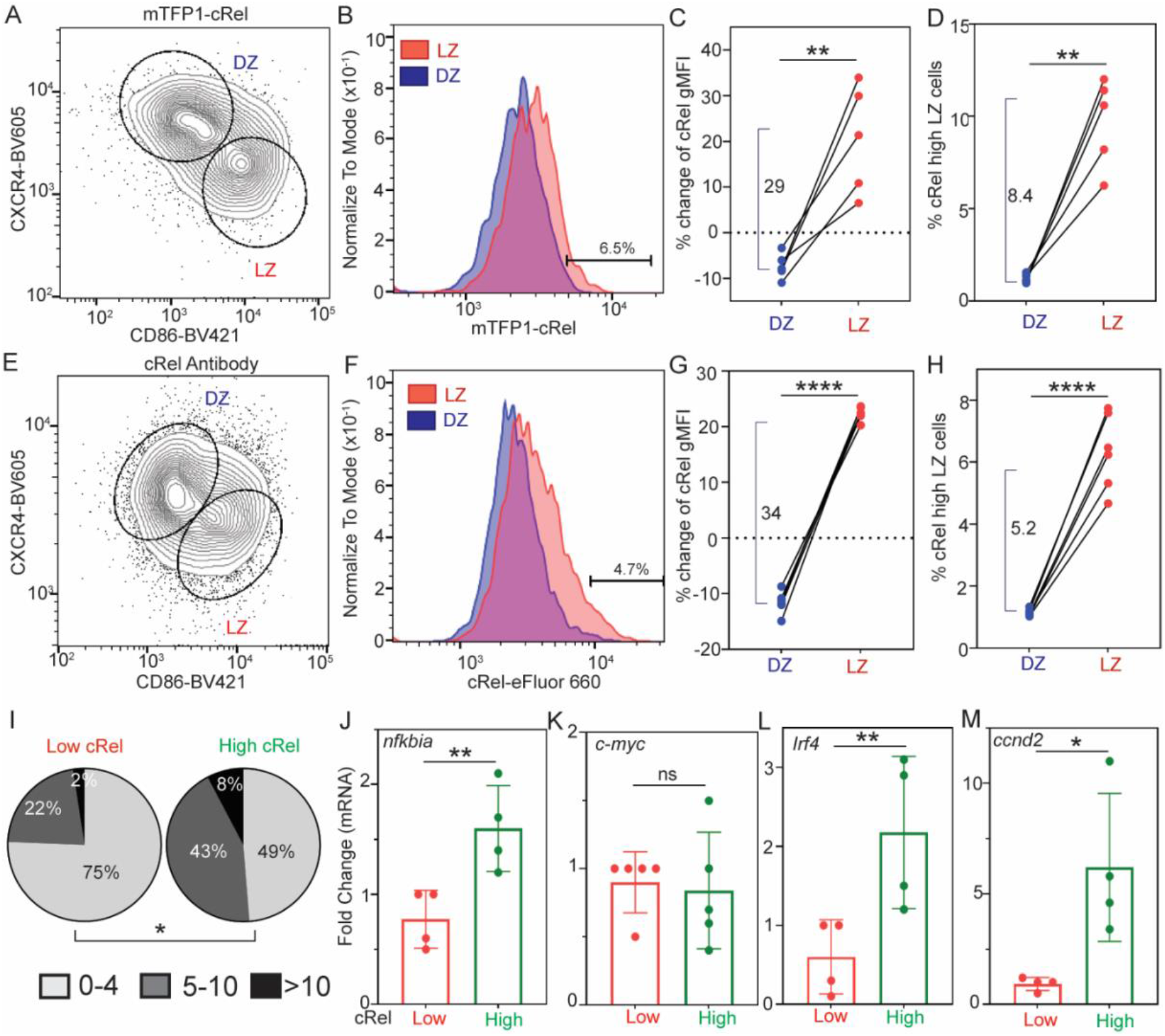
Increased cRel expression in LZ cells compared to DZ cells. WT and cRel^mTFP1^ mice were immunized, and cRel expression was measured in LZ cells (B220+CD95+GL7+CXCR4^low^CD86^high^) and DZ cells (B220+CD95+GL7+CXCR4^high^CD86^low^). **(A)** Flow cytometry plot of LZ cells and DZ cells in cRel^mTFP1^ mice. **(B)** Histogram of cRel expression in LZ cells and DZ cells in cRel^mTFP1^ mice. 6.5% is the proportion of LZ cells that show increased cRel expression compared to DZ cells. **(C)** Proportion changes of cRel gMFI in LZ and DZ cells from GC B cells in cRel^mTFP1^ mice. gMFI represents cRel expression. 29 is the mean % increase in cRel expression in LZ cells compared to DZ cells. The proportion change of cRel expression is calculated as follows: 100x(X-Y)/Y. X represents the gMFI of LZ cells or DZ cells, and Y represents the gMFI of GC B cells, which includes both LZ and DZ cells. **(D)** Proportion of cRel high-expressing LZ and DZ cells in cRel^mTFP1^ mice. 8.4 is the mean % increase in cRel high-expressing cells in LZ cells compared to DZ cells. **(E)** Flow cytometry plot of LZ cells and DZ cells in WT mice stained with cRel antibody. **(F)** Histogram of cRel expression in LZ cells and DZ cells in WT mice. 4.7% is the proportion of LZ cells that show increased cRel expression compared to DZ cells. **(G)** Proportion changes of cRel gMFI in LZ and DZ cells from GC B cells in WT mice. gMFI represents cRel expression. 34 is the mean % increase in cRel expression in LZ cells compared to DZ cells. **(H)** Proportion of cRel high-expressing LZ and DZ cells in WT mice. 5.2 is the mean % increase in cRel high-expressing cells in LZ cells compared to DZ cells. **(I)** Frequency of mismatch errors in the JH558 intronic sequence spanning 700 bp in low and high cRel-expressing LZ cells. The proportions of cells with 0-4 (light gray), 5-9 (gray), and more than 10 (black) mismatches are shown in the respective color zone. The number of clones sequenced in low and high cRel LZ cells is 41 and 39, respectively. Fisher’s exact test is used to compare the number of mutations between groups. Three independent experiments were performed, and 2-4 mice were pooled together for one experiment. **(J-M)** Quantitative measurement of mRNA expression of *nfkbia* **(J)**, *c-myc* **(K)**, *Irf4* **(L)**, and *Ccnd2* **(M)** by qPCR in low and high cRel LZ cells. Ubiquitin C (UBC) is used as a housekeeping control. For **(C), (D), (G)**, and **(H)**, one dot represents one mouse; for **(J-M)**, one dot represents one replicate and 3-5 mice pooled per replicate. Paired student’s t-test was performed. ns: not significant, *p < 0.05, **p < 0.01, ***p < 0.001, and ****p < 0.0001.

We investigated whether increased cRel expression is associated with enhanced SHM. We isolated LZ cells that express high or low cRel (Fig. S2A) and sequenced DNA from the JH4 intronic enhancer of IgH, which lies downstream of the V_J558_DJ_H4_ element (28, 29). We analyzed the frequency of mismatch errors in the 700-bp intronic sequence of JH558 in LZ cells expressing low or high cRel. We categorized mismatches into three groups: 0-4, 5-9, and >10 as described by Hanson *et al*. (28). For cRel low LZ cells, 75% of cells showed 0-4 mismatches, 22% showed 5-9 mismatches, and 2% showed 0-4 mismatches (Fig. 2I). For cRel high LZ cells, 49% of cells showed 0-4 mismatches, 43% showed 5-9 mismatches, and 8% showed 0-4 mismatches (Fig. 2I). Hence, cRel high LZ cells showed an increased number of mutations compared to cRel low LZ cells, suggesting cRel high LZ cells were likely positively selected.

*nfkbia* is a well-established NFkB target gene, validated across multiple cell types (30, 31). To confirm that cRel high LZ cells enhance *nfkbia* expression, we examined the expression of *nfkbia*. We found that cRel high LZ cells had enhanced expression of *nfkbia* relative to cRel low LZ cells, suggesting cRel high LZ cells have increased NFκB activity (Fig. 2J and S2B). Recent studies have shown that positively selected LZ cells primarily have two fates: either enter the DZ for an additional round of SHM to improve antibody affinity or generate pre-PC and exit GC to become mature PCs (14). Previous studies have shown that high cMyc-expressing LZ cells preferentially enter the DZ (4, 11, 12). To assess whether enhanced SHM in cRel high LZ cells correlates with increased cMyc expression, we measured cMyc expression at the mRNA level. We found that *cmyc* expression remained similar between cRel high and low LZ cells, which was confirmed by using two distinct genes (*ubc* and *β-actin*) as controls, suggesting cRel high LZ cells do not preferentially enter the DZ (Fig. 2K and S2C). *cmyc* is a low-expressed gene; hence, no difference in *cmyc* between cRel low and high could be attributed to the low level of *cmyc* expression (32, 33). Previous studies have shown that pre-PCs express higher levels of IRF4 and cyclin D2 (6, 14, 34, 35). To determine whether cRel high LZ cells have enhanced IRF4 and high cyclin D2 expression compared to cRel low LZ cells, we measured *irf4* and *ccnd2* (gene encoding cyclin D2) expression in cRel high and low LZ cells. We found increased *irf4* and *ccnd2* expression in cRel high LZ cells relative to cRel low LZ cells, which was confirmed by using *ubc* and *β-actin* as controls, suggesting cRel high LZ cells are likely enriched for higher pre-PC relative to cRel low LZ cells (Fig. 2L-M, and Fig. S2D-E). In conclusion, we show that cRel high LZ cells exhibit increased SHM and higher expression of IRF4 and cyclin D2 than cRel low LZ cells.

### High cRel-expressing light-zone GC B cells show an increased proportion of pre-PCs

To determine whether cRel high LZ cells are preferentially enriched for pre-PC, we performed single-cell (sc) RNA profiling of cRel high and low LZ cells. scRNA sequencing (scRNA-seq) was performed on ∼851 cRel high LZ cells and ∼1164 cRel low LZ cells using the 10X Genomics platform. We found that cRel high LZ cells are enriched for NFκB target gene signature (Fig. S2F) and increase expression of *nfkbia, irf4*, and *ccnd2* in cRel low LZ cells, whereas *cmyc* expression remained similar between the two groups (Fig. S2G-J). Hence, RNA-seq analysis confirms the increased expression of *nfkbia, irf4*, and *ccnd2* as detected by qPCR (Fig. 2 and Fig. S2). We found that cRel high and low LZ cells were transcriptionally different, and pathway analysis shows differential enrichment of cellular and signaling pathways in cRel high and low LZ cells (Fig. S3A-C). cRel high LZ cells were overall enriched for B cell receptor signaling and cytokine response, while cRel low LZ cells were enriched for MHC I response pathways as well as lymphoid progenitor cell differentiation. BCR signaling in GC B cells enhances enrichment of the NFκB pathway and the survival of GC B cells (36). Chronic BCR signaling in mature B cells lead to activation of cRel, and cRel is required for the survival of mature B cells (37, 38). A previous study shows that CD40 signaling activates NFκB cRel in GC B cells (13). Enrichment of the BCR signaling pathway in cRel high LZ cells likely suggests that the combination of BCR and CD40 signaling induces high cRel expression, whereas CD40 signaling alone induces low cRel expression in GC B cells.

We performed dimensional reduction using the Uniform Manifold Approximation and Projection (UMAP) algorithm on the scRNA-seq data. The analysis of the scRNA-seq dataset for both the cRel high and low LZ cells identified 5 clusters, which were annotated based on their gene expression signatures (Fig. 3A). We found that cluster 1 showed higher expressions of *irf4, ccnd2, myc*, and lower expression of *bcl6*, suggesting a likely pre-PC cluster (Fig. 3B-E) (39). Previous studies have shown that pre-MBCs enhance expression of cell-survival gene *bcl2*, exhibit increased expression of cell surface receptor *ccr6*, and increased expression of the transcription regulator *hhex* (7, 9, 36). Cluster 0 cells showed higher expression of *ccr6, hhex*, and *bcl2* suggesting a likely pre-MBC cluster (Fig. 3F-H). Interestingly, we found that maximum enrichment of NFκB target gene signature in the pre-PC cluster (Fig.3I). The analysis of cRel high and low LZ cells showed that cRel high LZ cells are enriched for pre-PC, pre-MBC, and G2-M phase cells relative to cRel low LZ cells, whereas cRel low LZ cells were enriched for G1 and S phase of the cell cycle (Fig. 3J-K and Fig. S3D-F). Interestingly, we found that a higher proportion of cRel high LZ cells in the pre-PC cluster expressed *irf4* than in cRel low LZ cells (Fig. 3L). A higher proportion of cRel high LZ cells in the pre-PC cluster expressed *ccnd2*, and also, the amplitude of *ccnd2* expression was higher than in cRel low LZ cells (Fig. 3L). Further, we found that *cmyc* was enriched only in the pre-PC cluster, and the amplitude of *cmyc* expression was higher in the pre-PC of cRel high LZ cells than in cRel low LZ cells (Fig. 3L). We found that *hhex* expression was higher, and a higher proportion of cells expressed *Bcl2* in the pre-MBC cluster of cRel high LZ cells compared to cRel low LZ cells (Fig. 3M). A similar proportion of cells expressed *ccr*6, and the amplitude of *ccr*6 expression remained similar in the pre-MBC cluster of cRel high and low LZ cells (Fig. 3M). Overall, sc RNA-seq analysis suggests potential enrichment of pre-PC and pre-MBC in cRel high LZ cells compared to cRel low LZ cells. Since *cmyc* remains similar between cRel low and high LZ cells, cRel high LZ cells are likely not enriched for the DZ reentry cells.

**Figure 3:**
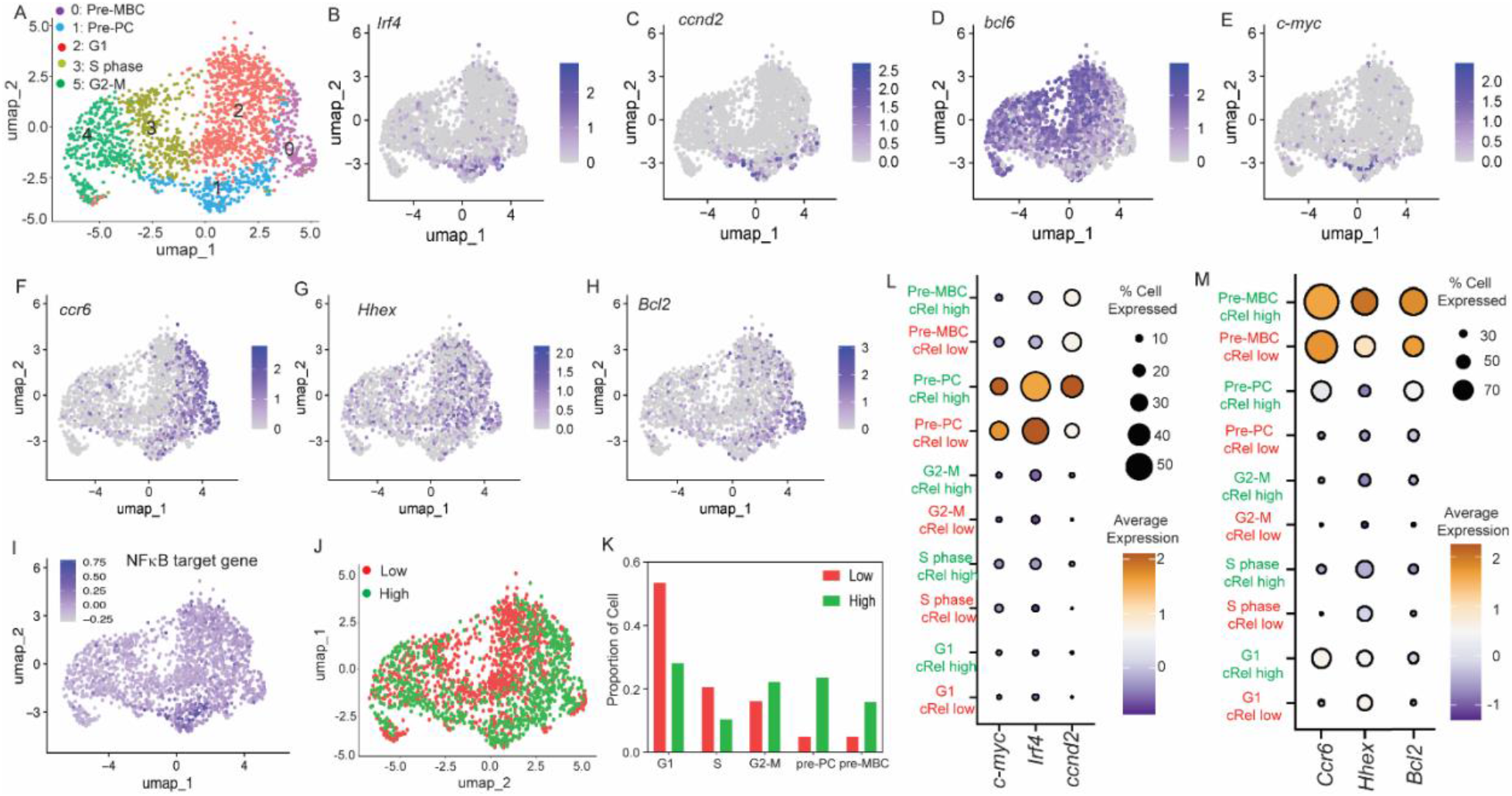
Single-cell RNASeq analysis project an increase in the proportion of pre-PCs and pre-MBCs in cRel high LZ cells. cRel low and high cRel LZ cells were flow sorted from cRel^mTFP1^ immunized mice for single-cell RNASeq. **(A)** UMAP plot projected 5 different cell clusters of all cell transcriptomes, which include cRel low and high LZ cells. These clusters are categorized by gene expression associated with cell cycle stages (G1, S, and G2-M phases) and by precursor MBC (pre-MBC) and precursor PC (pre-PC) populations. **(B-I)**. UMAP plot showing gene expression of selected genes relevant to pre-PC, *Irf4* **(B)**, *Ccnd2* **(C)**, *Bcl6* **(D)**, and *c-myc* **(E)**, and relevant to pre-MBC *ccr6* **(F)**, *Hhex* **(G)**, and *Bcl2* **(H)**, and NFκB target genes **(I)** (see materials and methods section for NFκB target gene list). The expression level is indicated by the scale bar shown on the side of each plot. The scRNA pipeline incorporates natural-log transformation using log1p. **(J)** Overlay of cRel low and high LZ cells on the UMAP plot. The number of cells sequenced in cRel low and high LZ cells is ∼1164 and ∼851, respectively. **(K)** Bar graph on the proportion of cells in each cluster in cRel low and high LZ cells. **(L)** Comparison of pre-PC relevant genes (*c-myc, Irf4*, and *Ccnd2*) between cRel low and high LZ cells in the G1, S-phase, and G2-M, pre-PC, and pre-MBC. The expression level is indicated by the scale bar on the side of each plot, and the proportion of cells expressing these genes is denoted by the circle size on the side of each plot. **(M)** Comparison of pre-MBC relevant genes (*Ccr6, Hhex*, and *Bcl2*) between cRel low and high LZ cells in the G1, S-phase, and G2-M, pre-PC, and pre-MBC. The expression level is indicated by the scale bar on the side of each plot, and the proportion of cells expressing these genes is denoted by the circle size on the side of each plot.

We investigated whether the potential enrichment of pre-PC and pre-MBC in cRel high LZ cells, as evident at the gene expression level, was confirmed at the phenotypic level using flow cytometry. We measured pre-PCs within LZ cells, defined by well-established surface and intracellular markers (B220+GL7+CD95+CXCR4^low^CD86^high^Bcl6^low^IRF4^high^CD69+) as previously described (6). We found that GC-derived pre-PC was higher in cRel high LZ cells than in cRel low LZ cells in cRel^mTFP1^ immunized mice (pre-PC in cRel high and low LZ cells: 11.6% and 1.9%, respectively), with a mean increase of 10% (Fig. 4A-C). To confirm that cRel high LZ cells have increased pre-PC compared to cRel low LZ cells, we measured pre-PC in cRel high LZ cells using intracellular cRel expression with cRel monoclonal antibody in WT mice. We found that cRel high LZ cells were enriched for pre-PC compared with cRel low LZ cells (pre-PC in cRel high and low LZ cells: 9.9% and 1%, respectively) with a mean increase of 9% (Fig. 4D-F, and Fig. S4A-E). Therefore, our study suggests that cRel high LZ cells are enriched for pre-PC. We measured pre-MBCs within GC B cells, defined by well-established surface markers (B220+GL7+CD95+CXCR4^low^CD86^high^CCR6+), as previously described (9). We found that GC-derived pre-MBC was slightly higher in cRel high LZ cells than in cRel low LZ cells in cRel^mTFP1^ immunized mice (pre-MBC in cRel high and low LZ cells: 15% and 12.5%, respectively), with a mean increase of 2.4% (Fig. 4G-I). On the contrary, we found that the proportion of pre-MBC in cRel high and low LZ cells remains very similar (pre-MBC in cRel high and low LZ cells: 16.9% and 17.3%, respectively) with a mean reduction of 1% using intracellular cRel antibody staining (Fig. 4J-L and Fig. S4F-G). The pre-MBC proportion in cRel high LZ cells showed a small increase (2.4%) using cRel^mTFP1^ immunized mice, whereas a minor decrease (1%) was observed with intracellular antibody staining. Hence, we conclude that pre-MBC remains similar between cRel high and low LZ cells. In conclusion, by combining sc RNA-seq and phenotyping (flow cytometry), we show that cRel high LZ cells are enriched for pre-PCs rather than pre-MBCs.

**Figure 4:**
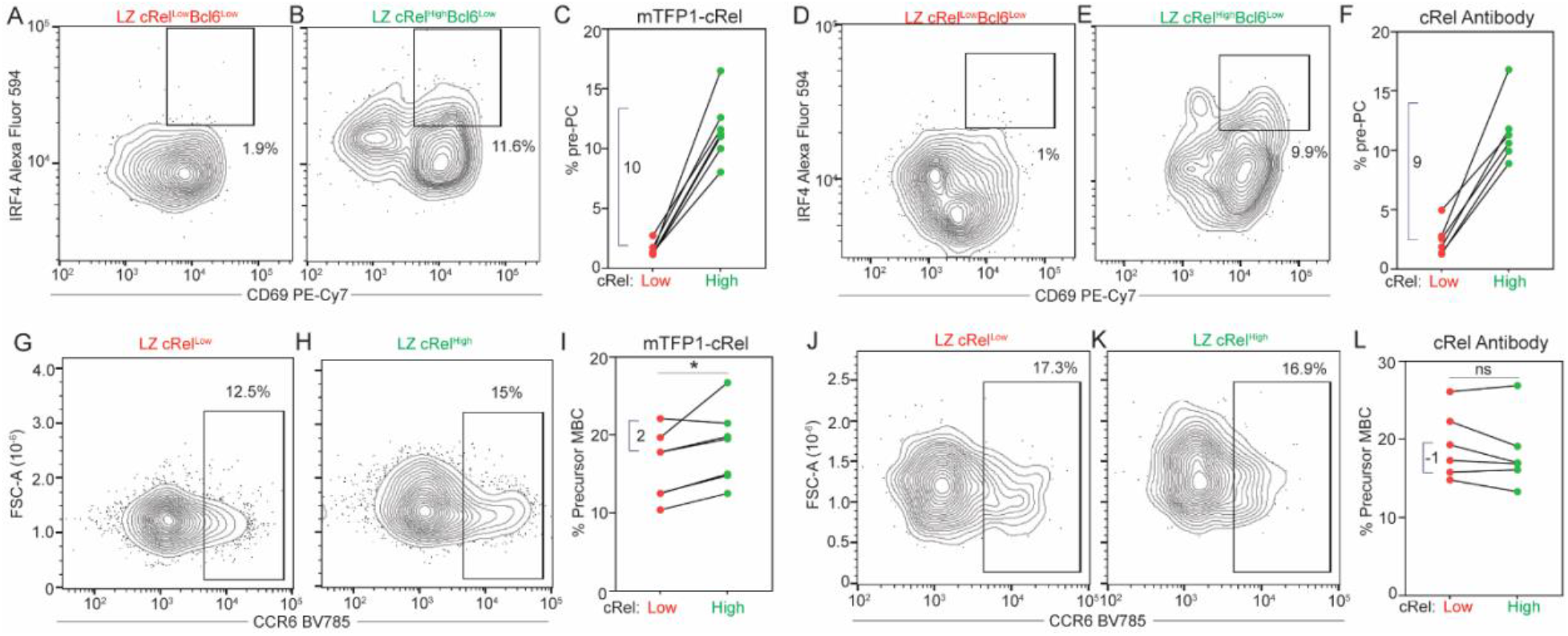
High cRel-expressing LZ cells are enriched for precursor PCs, not pre-MBCs. WT and cRel^mTFP1^ mice were immunized, and enrichment of pre-PCs and pre-MBCs was measured in low and high-cRel-expressing LZ cells. **(A-B)** Flow cytometry plot of pre-PC (B220+GL7+CD95+CXCR4^low^CD86^high^Bcl6^low^IRF4^high^CD69+) in cRel low **(A)** and high **(B)** LZ cells in cRel^mTFP1^ mice. 1.9% and 11.6% are the proportions of pre-PC in low and high cRel LZ cells, respectively. **(C)** Proportion of pre-PC in low and high cRel high-expressing LZ in cRel^mTFP1^ mice. 10 is the mean % increase of pre-PC population in cRel high-expressing LZ cells compared to cRel low-expressing LZ cells. **(D-E)** Flow cytometry plot of pre-PCs in cRel low **(D)** and high **(E)** LZ cells in WT mice stained with cRel antibody. 1% and 9.9% are the proportions of pre-PC in low and high cRel LZ cells, respectively. **(F)** Proportion of pre-PC in low and high cRel high-expressing LZ in WT mice. 9 is the mean % increase of pre-PC population in cRel high-expressing LZ cells compared to cRel low-expressing LZ cells. **(G-H)** Flow cytometry plot of pre-MBCs (B220+GL7+CD95+CXCR4^low^CD86^high^CCR6+) in cRel low **(G)** and high **(H)** LZ cells in cRel^mTFP1^ mice. 12.5% and 15% are the proportions of pre-MBC in low and high cRel LZ cells, respectively. **(I)** Proportion of pre-MBC in low and high cRel high-expressing LZ in cRel^mTFP1^ mice. 2 is the mean % increase of pre-MBC population in cRel high-expressing LZ cells compared to cRel low-expressing LZ cells. **(J-K)** Flow cytometry plot of pre-MBCs in cRel low **(J)** and high **(K)** LZ cells in WT mice stained with cRel antibody. 17.3% and 16.9% are the proportions of pre-MBC in low and high cRel LZ cells, respectively. **(L)** Proportion of pre-MBC in low and high cRel high-expressing LZ in WT mice. -1 is the mean % decrease of pre-MBC population in cRel high-expressing LZ cells compared to cRel low-expressing LZ cells. One dot represents one mouse. Paired Student’s t-test was performed. ns: not significant, *p < 0.05, **p < 0.01, ***p < 0.001, and ****p < 0.0001.

In this work, we investigated the function of cRel in GC B cell output at physiologic expression levels using cRel fluorescence reporter mice. We found that cRel expression is higher in GC B cells than in non-GC B cells, including MBCs (Fig. 1). A previous study shows that cRel-deleted GC B cells exhibit reduced SHM, whereas another study shows that cRel overexpression does not enhance SHM (19, 23). We have shown that at the physiologic expression levels, high cRel LZ cells exhibit increased SHM (Fig. 2I); thus, our study resolves the conflicting functions of cRel in SHM observed in cRel-deleted and cRel-overexpressed B cells. We found that cRel high LZ cells express higher IRF4 and cyclin D2, which are key in generating pre-PCs. cMyc expression remains similar between cRel high and low LZ cells (Fig. 2 and S2), suggesting that at physiologic expression levels, low cRel expression is sufficient to induce the gene expression program for reentry to DZ. Single-cell gene expression and flow cytometry-based phenotyping show enrichment for pre-PCs rather than for pre-MBCs (Fig. 3 and 4). Thus, we conclude that cRel high LZ cells favor enrichment of pre-PCs. Previously, we have shown that cRel regulates a complex gene regulatory network of B cell differentiation to PC by inducing IRF4, which in turn induces Blimp1, and cRel also inhibits Blimp1 by inducing Bach2, which represses Blimp1 (27). cRel plays a complex role in controlling gene regulatory networks and the differentiation of GC B cells (2, 40). While cRel-deleted and cRel-overexpressing GC B cells revealed that cRel is key to GC B cell formation and maintenance, the complex role of cRel in controlling the gene regulatory network within GC B cells cannot be fully elucidated by gene-knockout or overexpression systems; it is revealed by natural variation in cRel expression in this study. In line with this, we recently demonstrated that the natural variation of cRel correlates with both activation and termination of the proliferation program in B cells, which was not captured in previous studies utilizing cRel knockout B cells (16, 17, 41). Positively selected LZ cells primarily differentiate into either DZ cells for proliferation or PCs (4). The proliferation of positively selected LZ into DZ cells is regulated by cMyc expression, and PC differentiation is regulated by the amplitude of both IRF4 and Myc expression in LZ cells (4, 39). Here we report that cRel expression correlates with SHM and regulates pre-PC gene expression in LZ cells. We propose that cRel acts as a switch that determines whether positively selected LZ cells differentiate into PCs rather than DZ cells.

## Materials and Methods

### Mice and immunization

Mice in this study were maintained under specific-pathogen-free conditions supervised by the University of Utah Institutional Animal Care and Use Committee. Wildtype C57BL/6J (referred here as WT) (Strain #:000664) were either purchased from Jackson Laboratory or breed in house. mTFP1-cRel (cRel^mTFP1^) mouse strain has been described previously (17). C57BL/6J and cRel^mTFP1^ mice were immunized intramuscularly in both hind limbs with 50 mg of NP-CGG (4-Hydroxy-3-nitrophenylacetyl Chicken Gamma Globulin) (Biosearch Technology#N-5055B5 and Creative Diagnostics#DAGB499) with AddaVax adjuvant (Invivogen#vac-adx-10) 1:1 ratio (42). Inguinal lymph nodes were collected at day 7 post-immunization for analysis. Both male and female mice were used in this study, with an age range of 8 to 14 weeks.

### Cell preparation and flow cytometry

Inguinal lymph nodes were harvested at day 7 post-immunization. Lymph nodes were disrupted by gently crushing with a syringe plunger in B cell media (RPMI-1640, 20 mM HEPES, 10% FBS (Omega scientific international# FB-11), 1x Penicillin-Streptomycin, 5 mM L-glutamine, 5 mM MEM non-essential amino acids, 1 mM Sodium Pyruvates, and 55 mM 2-mercaptoethanol) and cell suspensions were filtered through a 40 mm sterile cell strainer (Fisher Scientific#22-363-547). Cells were centrifuged and resuspended in flow buffer (PBS, 2.5% FBS, 2 mM EDTA, and 0.02% sodium azide). 2×10^6^ cells were incubated with Fc receptor blocker anti-CD16/32 (Clone: 2.4G2) for 10 min on ice before staining with fluorochrome-conjugated antibodies for cell surface staining: anti-B220 (clone: RA3-6B2, anti-CD19 (clone: 1D3/CD19), anti-CD95 (clone: Jo2), anti-GL7 (clone: GL7), anti-CXCR4 (clone: L276F12), anti-CD86 (clone: GL-1), anti-CD38 (clone: 90), anti-IgD (clone: 11-26c.2a), anti-CD3e (clone KT3.1.1), and anti-NK1.1 (clone: DX5). Fluorochrome-conjugated cell-surface antibodies were incubated for 30 min on ice, with tapping every 10 minutes. Cells were centrifuged and resuspended in 250 mL flow buffer. Cells were filtered through a 40 mm cell strainer and incubated for 5-10 min with 5 mL of viability dye 7-AAD (7-amino-actinomycin D) (Biolegend# 420404). Cells were acquired using flow cytometry with a Cytek Aurora or a CytoFlex (Beckman Coulter).

For cRel intracellular staining, 4-5×10^6^ cells were incubated with Zombie NIR Fixable Viability dye (Biolegend# 423105) for viability staining at room temperature (RT) for 30 min in PBS. Cells were washed and resuspended in flow buffer. Cells were incubated with an Fc receptor blocker, then stained for cell surface markers as described above. cRel intracellular staining was performed using the Fix and Perm cell permeabilization kit (Thermo Fisher Scientific #GAS003), which stains both nuclear and cytoplasmic proteins, to assess cRel levels in GC B cells and MBCs, as described by the manufacturer’s protocol. Briefly, cells were fixed with 100 mL of reagent A at RT for 15 min, then washed with PBS. Cells were incubated with 100 mL of reagent B with anti-cRel (clone: 1RELAH5) at RT for 20 min, then washed twice with PBS.

10-15×10^6^ cells were used for intracellular staining of Bcl6, IRF4, and cRel to detect pre-PC and pre-MBC. Cell viability (Zombie NIR Fixable Viability) and cell surface staining are described above. This flow panel contains 4 or more antibodies conjugated with violet fluorochromes, and intracellular staining was performed with the Foxp3/Transcription Factor Staining kit (eBioscience #00-5523-00). BD Horizon Brilliant staining buffer (BD Biosciences #563794) was used for cell-surface staining for this flow panel to achieve good separation between negative and positive populations. Intracellular staining with Foxp3/Transcription Factor Staining kit was performed according to the manufacturer’s protocol. Briefly, cells were fixed with 600 mL of fixation buffer at RT for 20 min, then washed twice with 200 mL permeabilization buffer. Cells were incubated with 300 mL of permeabilization buffer with anti-Bcl6 (clone: K112-91), anti-IRF4 (clone: IRF4.3E4), and/ or anti-cRel (clone: 1RELAH5) at RT for 20 min, then washed twice with PBS. Bottom and top 40% cRel low and high LZ cells were gated, and then the proportion of the pre-PCs and pre-MBCs was measured in the cRel low and high LZ cells.

Cells were analyzed in FlowJo V10, and dead cells were excluded. Gating strategies are as follows: GC B cells B220+CD95+GL7+ (or B220+CD19+CD3e-NK1.1-IgD^low^CD38-GL7+), non-GC B cells B220+CD95-GL7-, MBCs (or B220+CD19+CD3e-NK1.1-IgD^low^CD38+GL7-), LZ cells B220+CD95+GL7+CD86^high^CXCR4^low^, DZ cells B220+CD95+GL7+CD86^low^CXCR4^high^, pre-PCs B220+CD95+GL7+CD86^high^CXCR4^low^Bcl6^low^IRF4^high^CD69+, and pre-MBC B220+CD95+GL7+CD86^high^CXCR4^low^CCR6+.

### cRel low and high LZ cells isolation

cRel^mTFP1^ mice were immunized, and 3-5 mice were pooled together for flow sorting of cRel low and high LZ cells. Inguinal Lymph nodes were isolated at day 7 post-immunization. 30-40×10^6^ cells in 1 mL were used for cell surface staining in flow buffer as described above. 20 mL of 7-AAD was added 10 minutes before flow sorting. Bottom and top 15% cRel low and high LZ cells were flow sorted in a 1.5 mL centrifuge tube. Cells were sorted with Cytek Aurora using a 70 mm nozzle in a cold (4-8℃) environment.

### Quantitative PCR

50,000-20,000 cRel low- and high-LZ cells were flow sorted in cold PBS. Then quantitative PCR was performed with the SYBR Green Fast Advanced Cells-to-Ct Kit (Thermo Fisher Scientific #A35380) according to the manufacturer’s protocol. Briefly, flow sorted cells were lysed immediately and, after lysis reaction, Cells to CT lysates were stored at -20◦C. Lysates were thawed, and the Reverse Transcription (RT) reactions were performed. Completed RT reactions were stored at -80◦C. Then quantitative PCR was performed with the RT reactions.

### BCR somatic mutation in JH558 intron

50,000 cRel low- and high-LZ cells were flow sorted in B cell media for BCR mutational analysis in the JH4 Intronic enhancer of IgH as described by Hanson et al., (28). Sorted cells were washed with PBS, snap-frozen in dry ice, and stored at -80℃. Cells were thawed, and DNA isolation was performed using a QIAamp DNA Micro kit (QIAGEN), and DNA was amplified using nested PCR. Nested PCR products were used for secondary amplification using new primers and Taq DNA polymerase (Fisher). The PCR product was ligated into the TOPO TA Cloning Kit for sequencing (Invitrogen) and transformed in E.Coli (NEB® 5-alpha Competent E. coli) (New England Biolabs# C2987H). Colonies were miniprepped for DNA isolation and sequenced using T3 Primer, 5’ CTGTGTTCCTTTGAAAGCTGG3’, and sequences aligned to the germline JH558 intronic sequence using standard nucleotide BLAST. Primers are as follows: Nested Forward 1, 5′-AGCCTGACATCTGAGGAC-3′; Nested Reverse 1, 5′-TCTGATCGGCCATCTTGACTC-3′; Nested Forward 2, 5′-CATCTGAGGACTCTGCGGTCT-3′; Nested Reverse 2, 5′-CTGTGTTCCTTTGAAAGCTGG-3′.

### Single Cell RNASeq

10,000 cRel low- and high-LZ cells were flow sorted in B cell media containing 20% FBS. Sorted cells were washed with PBS and resuspended in 20 mL of 0.04% BSA for the 10X Genomics single cell RNA sequencing library. The sequencing library was prepared with the 10X Genomics Next GEM Single Cell 3’ Gene Expression kit v3.1. The resulting libraries were pooled and sequenced with NovaSeq Reagent Kit v1.5_150×150 bp Sequencing (∼400×10^6^ read-pairs were sequenced per condition). Libraries were sequenced to a read depth of at least 200,000 reads per cell with at least 88% sequencing saturation.

### Single Cell RNASeq and UMAP analysis

Post-sequencing, FASTQ files were generated and aligned to 10X Genomics’ mouse reference genome refdata-gex-mm10-2020-A with CellRanger version 7.0.0. The output HDF5 (Hierarchical Data Format version 5) cell barcode and gene UMI (unique molecular identifier) count matrix was then loaded into R version 4.3.1 and analyzed with the Seurat R package (version 5.2). Percentages of mitochondrial and ribosomal genes were calculated, and cells were clustered using 8000 variable genes with mitochondrial, ribosomal, and hemoglobin genes removed. Variable genes were scaled and used as input to compute 100 principal components. Of these, 60 principal components were used as input to UMAP for visualization and clustering. To calculate the cell cycle phase, Seurat’s Cell cycle scoring function was used, with human S-phase and G2M genes converted to their mouse equivalents.

The final NFκB target genes presented here were selected based on the well-established NFκB target genes relevant to B cell biology. We used Chen *et al*. and Zhao *et al*. to curate the NFκB target genes list (36, 43). These are NFκB genes selected in this paper. Rel, Rela, Relb, Nfkb1, Nfkb2, Nfkbia, Nfkbib, Nfkbiz, Bcl3, Bcl2, Tnfaip3, Bcl2a1a, Bcl2l1, Cd40, Cd80, Cd86, Icam1, Ccl3, Ccl5, Il6, Prdm1, Irf4, and myc.

## Author Contribution

KR conceptualized and designed the study. SR, KR, TT, BM, AK, and EVW performed research. SK performed preliminary experiments that contributed to develop the study. KR, SR, TT, and JD analyzed the data. KR wrote the paper with support from SR and TT. SR wrote the Materials and Methods section of the experimental section. TT wrote the method sections for the bioinformatics analysis and the figure legends for the single-cell analysis. All authors edited and approved the paper and are responsible for their data.

## Acknowledgment

We thank Jason Cyster for critically reading and providing feedback on this study. We thank the current and former lab members of the Roy lab for their critical discussion and feedback on this study. KR acknowledges funding from the National Institute of Allergy and Infectious Diseases (R56AI177789 and R01AI186945). We are thankful to the University of Utah Flow and Sequencing Core Facility. KR acknowledges assistance from Grammarly for grammar, punctuation, and paraphrasing, while retaining accountability for the content of the writing.

---

**Figure S1:**
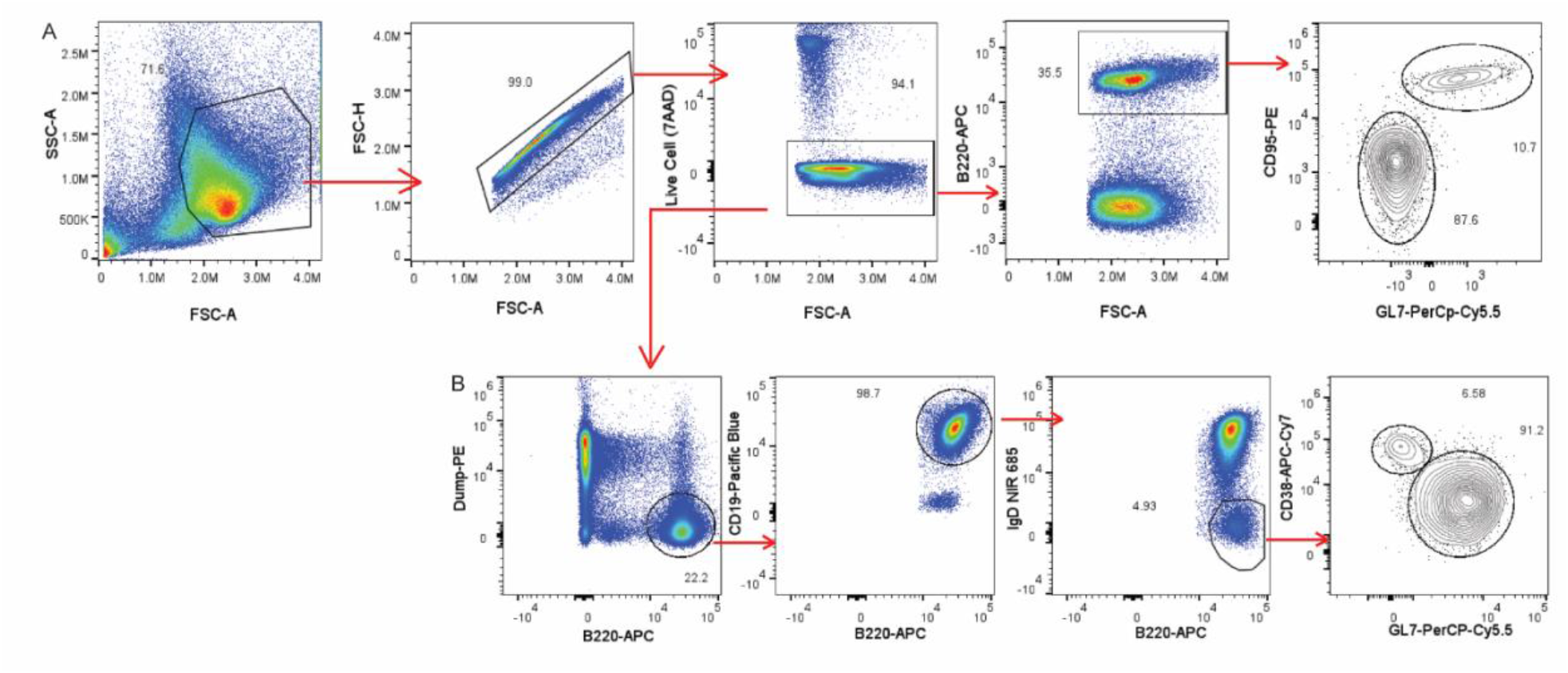
Gating strategies. **(A)** Gating strategy for GC B cells (B220+CD95+GL7+) and non-GC B cells (B220+CD95-GL7-) for Figure 1A-B. **(B)** Gating strategy for GC B cells (B220+CD19+CD3e-NK1.1-IgD^low^CD38-GL7+) and MBCs (B220+CD19+CD3e-NK1.1-IgD^low^CD38+GL7-) for Figure 1I and 1L.

**Figure S2:**
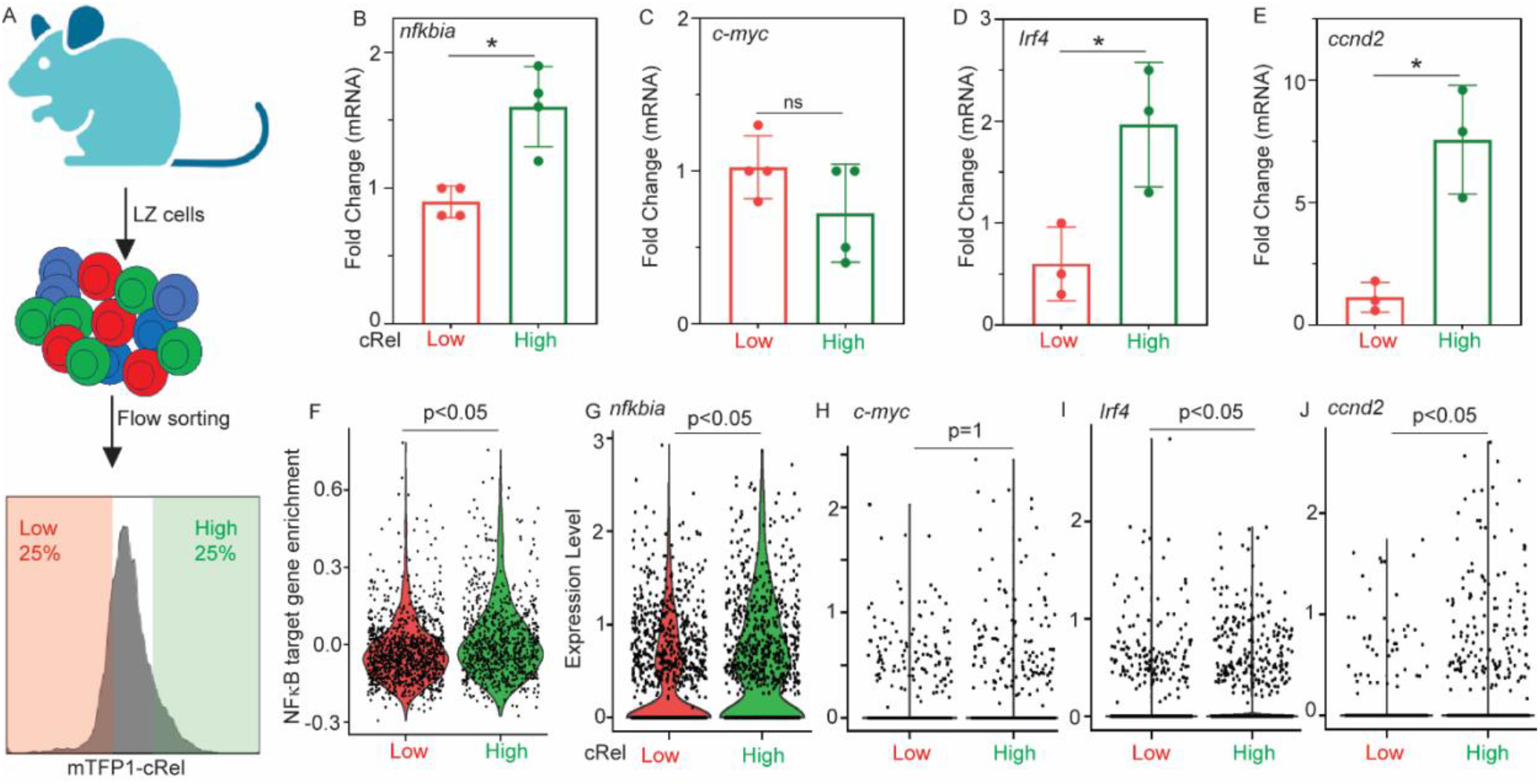
Expression of key regulators of GC B cells in cRel low and high LZ cells: **(A)** Schematic of experimental design of cRel low and high LZ cells isolation by flow sorting. cRel^mTFP1^ mice were immunized, and cRel low (bottom 25% in the histogram) and high (top 25% in the histogram) LZ cells were flow sorted. **(B-E)** Quantitative measurement of mRNA expression of *nfkbia* **(B)**, *c-myc* **(C)**, *Irf4* **(D)**, and *Ccnd2* **(E)** by qPCR in low and high cRel LZ cells. β-actin is used as a housekeeping control. **(F-J)** Violin plot depicting enrichment of NFκB target genes **(F)**, and expression of *nfkbia* **(G)**, *c-myc* **(H)**, *Irf4* **(I)**, and *Ccnd2* **(J)** by scRNASeq in low and high cRel LZ cells. For figure **(B-E)**, unpaired student’s t-test was performed. ns: not significant, *p < 0.05, **p < 0.01, ***p < 0.001, and ****p < 0.0001. For figure **(F-J)**, A Wilcoxon rank-sum test with a Bonferroni correction was performed to assess overall transcriptional changes in scRNA datasets comparing the cRel high LZ cell population with the cRel low LZ cells. NFκB target genes, *Nfkbia, Irf4*, and *Ccnd2* were significantly elevated in the cRel high LZ cell population.

**Figure S3:**
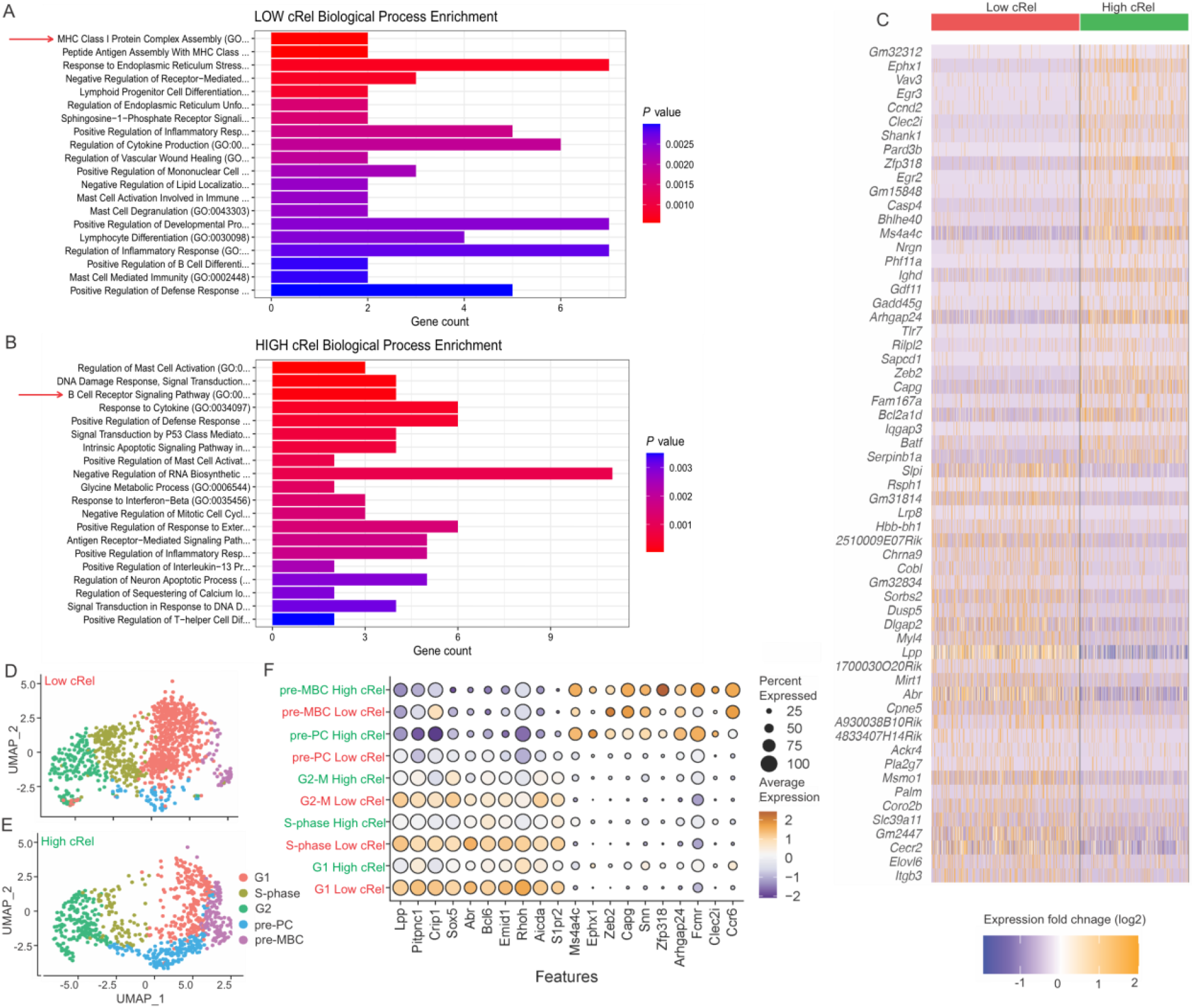
Low and high cRel-expressing LZ cells are transcriptionally different. Populations of cells expressing high and low levels of cRel show broadly different transcriptional profiles. **(A-B)** Pathway analysis of the top 150 differentially expressed genes between cRel low **(A)** and high **(B)** LZ cells using EnrichR against the 2025 Gene Ontology Biological Processes database. High cRel cells are overall enriched for B cell receptor signaling and cytokine response (indicated with red arrow), while low cRel samples are enriched for MHC I response pathways as well as lymphoid progenitor cell differentiation (indicated with red arrow). **(C)** The top 30 differentially expressed genes overall for high cRel cells and low cRel cells. **(D-E)** Shows distinct clustering patterns due to transcriptional differences, with low cRel LZ cells being enriched in the G1 and S-phase subclusters **(D)**, while high cRel LZ cells profiles are enriched in pre-PC, pre-MBC, and G2-M subclusters **(E). (F)** Within subcluster comparisons, identifying the top differentially expressed genes per subcluster for high cRel cells compared to low cRel cells.

**Figure S4:**
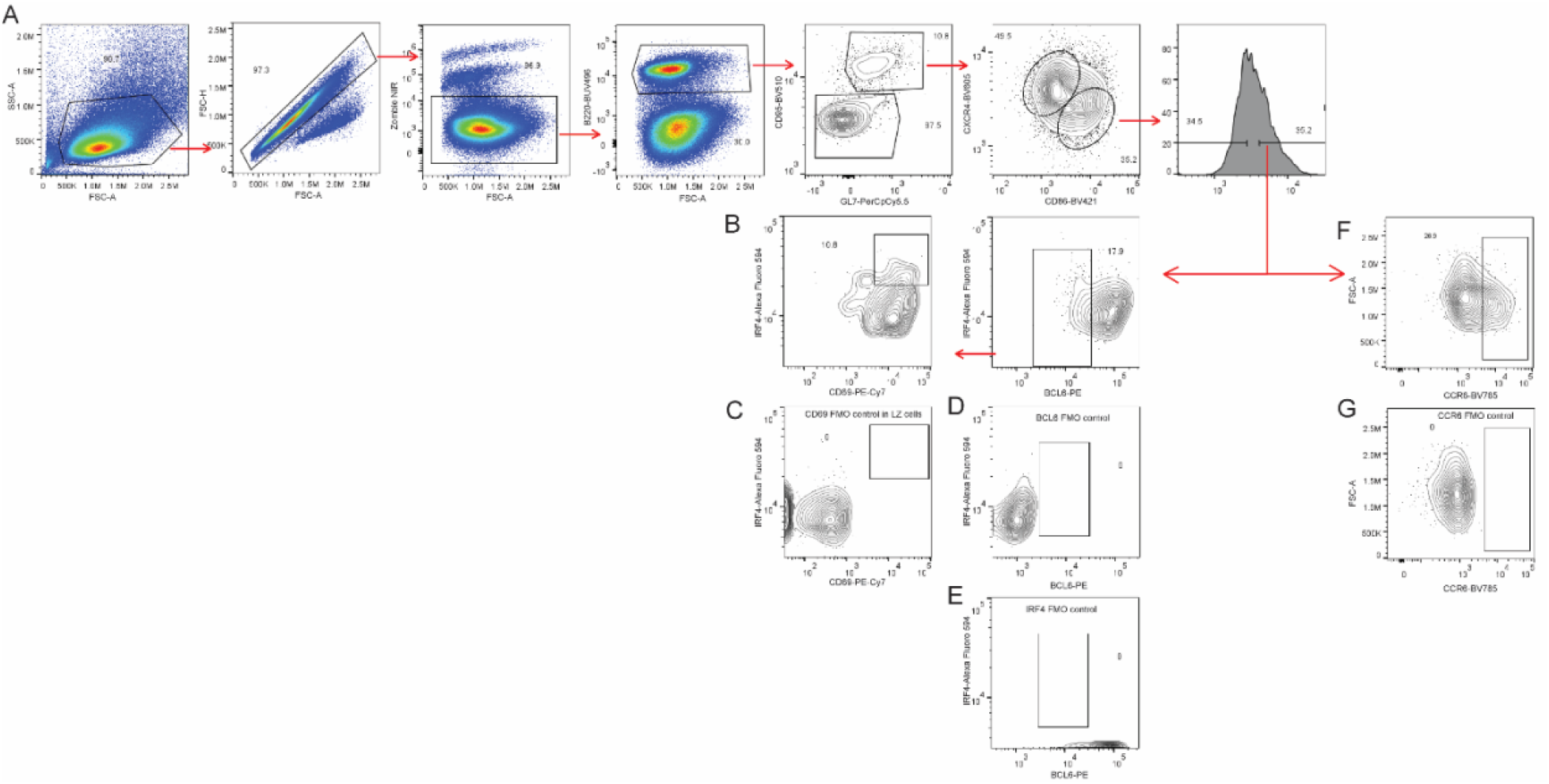
Gating strategy for pre-PCs and pre-MBCs. **(A)** Gating strategy of cRel low (bottom 30-40%) and high (top 30-40%) LZ cells. **(B)** Gating strategy of pre-PCs (B220+GL7+CD95+CXCR4^low^CD86^high^Bcl6^low^IRF4^high^CD69+) within cRel high LZ cells. A similar gating strategy is applied for cRel low LZ cells. **(C)** Fluorescence minus one (FMO) control for CD69 in LZ cell population (B220+GL7+CD95+CXCR4^low^CD86^high^Bcl6^low^IRF4^high^). **(D)** Fluorescence minus one (FMO) control for BCL6 in LZ cell population (B220+GL7+CD95+CXCR4^low^CD86^high^IRF4^high^CD69+). **(E)** Fluorescence minus one (FMO) control for IRF4 in LZ cell population (B220+GL7+CD95+CXCR4^low^CD86^high^Bcl6^low^CD69+). **(F)** Gating strategy of pre-MBCs (B220+GL7+CD95+CXCR4^low^CD86^high^CCR6+) within cRel high LZ cells. A similar gating strategy is applied for cRel low LZ cells. **(G)** Fluorescence minus one (FMO) control for CCR6 in LZ cell population (B220+GL7+CD95+CXCR4^low^CD86^high^).

